# plotsr: Visualising structural similarities and rearrangements between multiple genomes

**DOI:** 10.1101/2022.01.24.477489

**Authors:** Manish Goel, Korbinian Schneeberger

## Abstract

**Summary:** Third-generation genomic technologies have led to a sharp increase in the number of high-quality genome assemblies. This allows the comparison of multiple assembled genomes of individual species and demands for new tools for visualising their structural properties. Here we present plotsr, an efficient tool to visualize structural similarities and rearrangements between multiple genomes. It can be used to compare genomes on chromosome level or to zoom in on any selected region. In addition, plotsr can augment the visualisation with regional identifiers (e.g. genes or genomic markers) or histogram tracks for continuous features (e.g. GC content or polymorphism density).

**Availability and implementation:** plotsr is implemented as a python package and uses the standard matplotlib library for plotting. It is freely available under the MIT license at GitHub (https://github.com/schneebergerlab/plotsr) and bioconda (https://anaconda.org/bioconda/plotsr).

**Contact:** Manish Goel, Korbinian Schneeberger

## Introduction

Third-generation sequencing technologies like PacBio high-fidelity sequencing (HiFi) and Oxford-nanopore sequencing (ONT) have revolutionised genome assembly (Koren and Phillippy, 2015). Together with methods like Hi-C, trio-binning or gamete-binning (that perform efficient phasing and scaffolding of the contigs) haplotype-resolved, chromosome-level assemblies can easily be generated (Zhang *et al*., 2019; Koren *et al*., 2018; Campoy *et al*., 2020). This has led to a sharp increase in the number of available high-quality genome assemblies. Chromosome-level assemblies allow for the identification of not only small genomic differences (like SNPs and indels) but also for finding large structural rearrangements (SRs) like inversions and translocations. They are therefore considered as the gold standard for the identification of genomic differences (Simpson and Pop, 2015).

An interesting observation can be made when comparing the different genomes of an individual species (intra-genome comparison). The overall structure of these genomes is usually highly conserved as the genomes need to exchange chromosome arms during sexual reproduction. This implies the presence of extensive amounts of syntenic regions between the genomes, where the exchange of chromosome arms can happen without affecting the overall structure of the genomes. During genome assembly comparison these syntenic regions (the “syntenic backbone”) can be identified. All remaining regions in the genomes are structural rearrangements by definition and can then be classified into duplications, translocations or inversions based on their orientation and location in the genomes. Recently, we used these principles to develop SyRI, a tool to identify genomic differences in whole-genome assemblies of the same species (Goel *et al*., 2019).

In this work, we present **plotsr** (**plot s**tructural **r**earrangements) a novel visualisation tool for genomic differences, which is based on the synteny concepts included in SyRI. Using the syntenic backbone as a guide, plotsr can visualize structural differences on a chromosome level, as well as zoom in on specific regions. plotsr is a simple-to-use yet flexible and powerful visualisation tool. It can be used to compare multiple haploid genomes as well as different haplotypes of individual polyploid genomes. In addition, plotsr can mark specific loci as well as plot histogram tracks to show distributions of genomic features along the chromosomes.

### Implementation

plotsr is a command-line tool. It requires the assembly sequences (in fasta format) and the synteny and SRs information between the assemblies in a pairwise manner as input. For example, to visualise genomes A, B and C in this order, plotsr requires the comparison of A *vs*. B and B *vs*. C. These can be generated using SyRI or can be provided in BEDPE format. Firstly, plotsr validates that the assemblies and structural information are consistent. Then, by using the pairwise synteny between genomes, it groups homologous chromosomes across the genomes and plots the syntenic regions as well as SRs between them. The output plot can be generated in pdf, png or svg format. In addition to the genomic similarities and differences, plotsr can include markers at specific loci (e.g. genes, TEs, or genomic markers) using BED files. plotsr can also plot histogram tracks showing the distribution of genomic features along the chromosomes (e.g. distribution of SNPs, sequencing reads etc). This provides a visual comparison between sequence features and structural properties of the chromosomes. To adjust these plots to specific needs, plotsr includes multiple parameters to control the visual properties (colour, size, spacing etc) of genomes, markers and tracks.

plotsr can also be used to zoom in on specific regions in any of the input genomes. For this, plotsr identifies the corresponding homologous regions in all other genomes. This is a non-trivial task as some regions might include multiple rearrangements that obfuscate the syntenic regions in the other genomes. The identification of all syntenic regions would require whole-genome alignments of all genomes against the genome of interest implying the need for an all-vs-all genome alignment as input. This is computationally prohibitive once tens of genomes are involved. Instead, plotsr overcomes this challenge by using the syntenic backbone between the genomes to zoom in on any given region. For this, plotsr iteratively selects the regions syntenic to the selected region using pairwise genome comparisons until all genomes are covered. It then filters the structural information to only plot information overlapping these homologous regions resulting in a zoomed-in view of the genomes. Markers and feature tracks are also filtered automatically to plot those overlapping with the homologous regions.

## Results

We demonstrated the applicability of plotsr by visualising structural rearrangements between eight divergent strains of the model plant *Arabidopsis thaliana* (The Arabidopsis Genome Initiative, 2000; Jiao and Schneeberger, 2020) (Figure 1). For this, we performed seven pairwise whole-genome alignments using minimap2 and then identified synteny and structural rearrangements using SyRI (Li, 2018; Goel *et al*., 2019). In addition, we obtained some general genome features for further labelling (including gene annotation, population-wide polymorphism data and centromere coordinates (Lamesch *et al*., 2012; Alonso-Blanco *et al*., 2016; Giraut *et al*., 2011)). The visualization by plotsr showed that the genomes were predominantly syntenic (grey alignments) except for some large inversions including a 2.5MB inversion on Chr3 in the genome of Sha and a 1.1MB inversion on Chr4 in Col-0, which we labelled using plotsr as “Inversion 1” and “Inversion 2”. Using the annotation tracks, the increased density of SRs within the centromere regions became readily apparent in the visualization. Moreover, we also plotted the genes and frequency of SNPs along the chromosomes, which revealed the known inverse relationship between gene and SNP density (The Arabidopsis Genome Initiative, 2000) as well as the depletion of genes in the centromeres.

**Figure 1:**
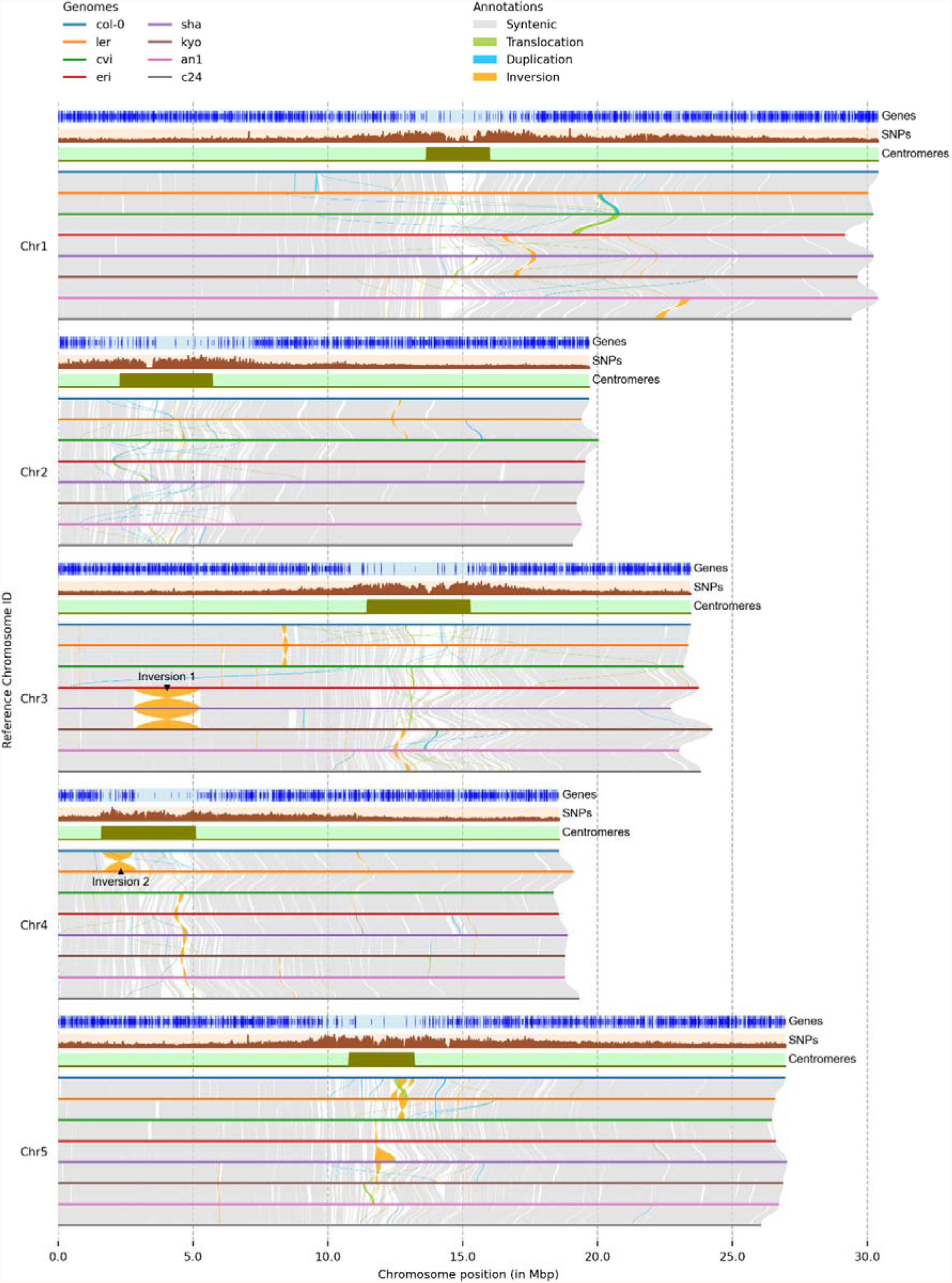
Example for a genome-wide visualization using plotsr. We used plotsr to visualize syntenic regions and structural rearrangements between eight strains of *A. thaliana*. The visualisation is created using plotsr without further modifications. It shows synteny and SRs between the eight genomes. Tracks for three genomic features: genes, number of SNPs and centromeric regions were included using optional parameters. In the genes track, small blue lines correspond to transcribed regions and long blues lines represent coding-sequences (CDS). We also marked two large inversions, one each on chromosome 3 and chromosome 4.

We also used the zoom-in option of plotsr to highlight the genomic differences at Chr3:6,600,000-10,000,000 in the genome of “col-0” (Figure 2). The region was provided as a command-line parameter to plotsr which then automatically filtered and plotted the syntenic regions in all of the remaining genomes. In this example, we observed two large regions without any alignments (labelled as “Not aligned 1” and “Not aligned 2”) that signify divergence between the two strains as well as two more inversions that become visible during zoom in.

**Figure 2:**
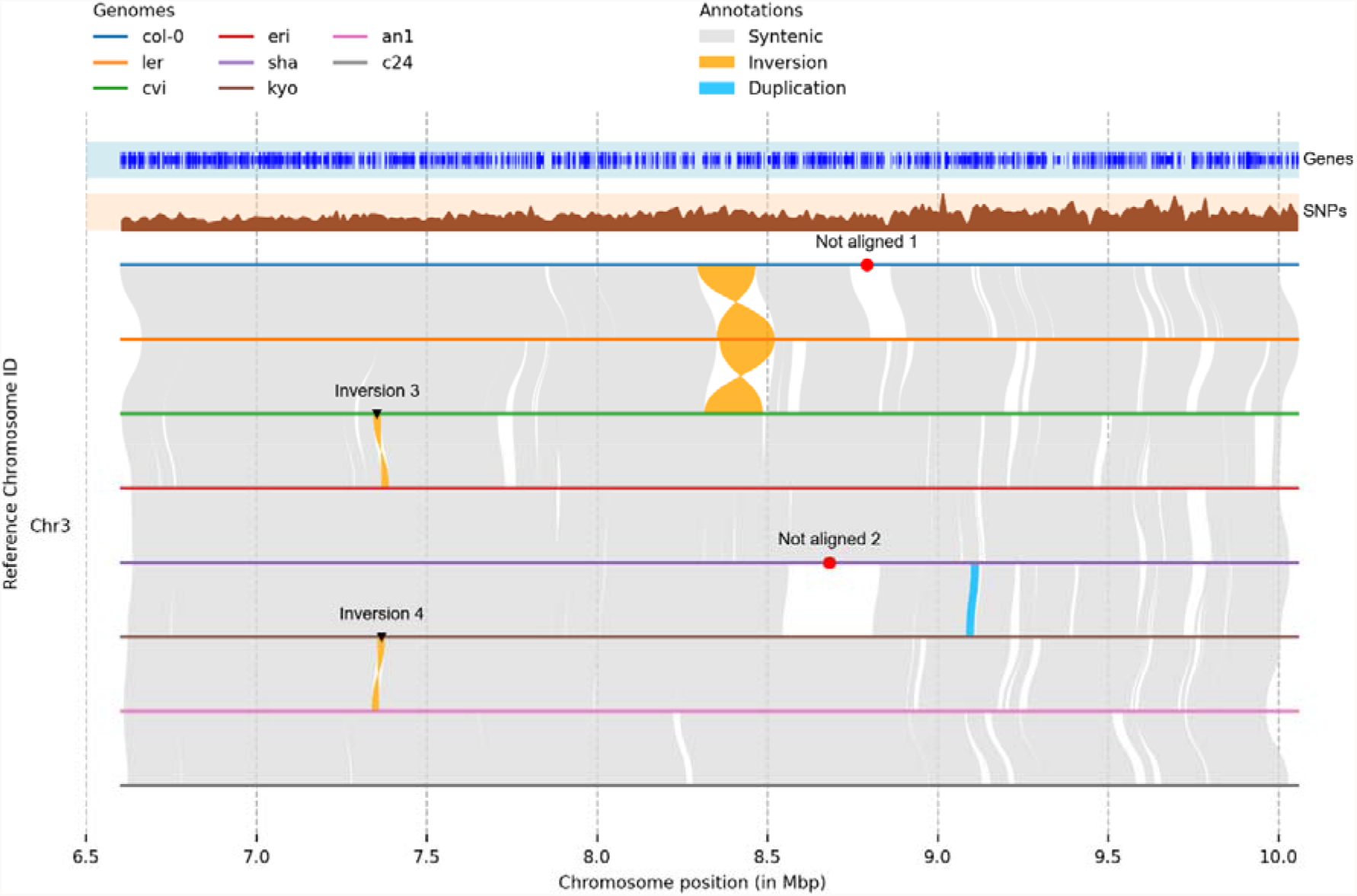
Using plotsr to zoom in on Chr3:6,600,000-10,000,000 in the “col-0” genome. The visualisation is created using plotsr without any further modifications. Zooming in on a specific location allows for resolved visualisation of the local genomic differences. Here, we observe two large not aligned regions and two smaller inversions, which have been labelled with plotsr.

## Conclusion

The advent of high-quality long-read sequencing technologies simplified the generation of high-quality genome assemblies. To support the visual analysis of such assemblies, we presented plotsr, a computational tool for visualising structural similarities and rearrangements between genomes. plotsr is highly adjustable and allows additional visualisation of genomic features as well as zoom-in views on specific regions. It generates publication-quality visualisations (Zamyatin *et al*., 2021; Zhang *et al*., 2021; Li *et al*., 2021) and will help towards a better understanding of differences within genomes. Given the great importance of genomic analysis in many research fields, we are continuously developing plotsr to add more useful parameters allowing for more control and customisation.

